# Understanding the link between functional profiles and intelligence through dimensionality reduction and graph analysis

**DOI:** 10.1101/2023.04.12.536421

**Authors:** F. Alberti, A. Menardi, D.S. Margulies, A. Vallesi

## Abstract

There is a growing interest in neuroscience for how individual-specific structural and functional features of the cortex relate to cognitive traits. This work builds on previous research which, using classical high-dimensional approaches, has proven that the interindividual variability of functional connectivity profiles reflects differences in fluid intelligence. To provide an additional perspective into this relationship, the present study uses a recent framework for investigating cortical organization: *functional gradients.* This approach places local connectivity profiles within a common low-dimensional space whose axes are functionally interretable dimensions. Specifically, this study uses a data-driven approach focussing on areas where FC variability is highest across individuals to model different facets of intelligence. For one of these loci, in the right ventral-lateral prefrontal cortex (vlPFC), we describe an association between fluid intelligence and relative functional distance from sensory and high-cognition systems. Furthermore, the topological properties of this region indicate that with decreasing functional affinity with the latter, its functional connections are more evenly distributed across all networks. Participating in multiple functional networks may reflect a better ability to coordinate sensory and high-order cognitive systems.

**Significant Statement:** The human brain is highly variable. In particular, the way brain regions communicate to one another – that is, how they are *functionally* connected – constitutes a neural fingerprint of the individual. In this study, we make use of a recent methodological approach to characterize the connectivity patterns of transmodal (closely linked to abstract processing) and unimodal (closely linked to sensory processing) brain regions in an attempt to explain how this balance affects intelligence. We show that the more the functional profile of executive control regions is distant to that of abstract processing, the better they are at integrating information coming from widespread neural systems, ultimately leading to better cognitive performance.

## 1. Introduction

Current neuroscience conceptualizes behavior as the result of the dynamic interaction of distributed communities of cortical regions and subcortical structures. Previous literature has examined these interactions in depth demonstrating that they possess certain stable topological features (1–3). However, the human brain also shows a high degree of structural and functional variability across individuals (4) owing to the interaction of genetic and environmental factors (5).

The unique functional and structural characteristics of an individual’s brain have been compared to a fingerprint, as they exhibit unique patterns of structural morphology (6), white matter tracts (7), and intrinsic functional connectivity (FC) (8) that can be used to accurately identify it. These different sources of interindividual variability have been mapped across the cortex demonstrating that association areas exhibit the most unique structural-functional patterns, making the dorsal-attention, frontoparietal (FPN), and default-mode (DMN) the most variable networks across individuals (4). Such variability of functional architecture reflects behavioral differences between subjects, and multiple studies have suggested that it is possible to predict behavioral measures from individual resting FC organization (4, 8). Existing evidence shows that functional topology is especially effective in predicting cognitive measures and, specifically, fluid intelligence, stressing the importance of associative regions and related networks for higher-order cognition (4). This work aims to provide additional insight into how FC variability in these regions translates into behavioral differences by adopting a low-dimensional perspective on functional organization.

With this aim, we adopt a framework that has been recently introduced to describe intrinsic FC patterns through a set of *functional dimensions* that reflect different organizational principles of brain activity (9). This approach exploits dimensionality reduction algorithms to reveal the principal components (latent dimensions) of FC data and blood oxygen level dependent (BOLD) time series (10). These dimensions are referred to as *functional gradients* and they describe smooth functional transitions along the cortical surface (Fig. 1). The first three gradients as they explain the majority of variance in the original data and recapitulate relatively clear functional axes. The principal gradient (explaining the most variance; Fig. 1A) is anchored in unimodal sensory-motor areas and peaks in transmodal association regions of the DMN; the second gradient (Fig. 1B) groups the visual network at one end and the somato-motor network at the opposite one; the third gradient (Fig. 1C) is anchored in the DMN and gradually progresses towards the FPN or multiple demand network (MD).

**Figure 1.**
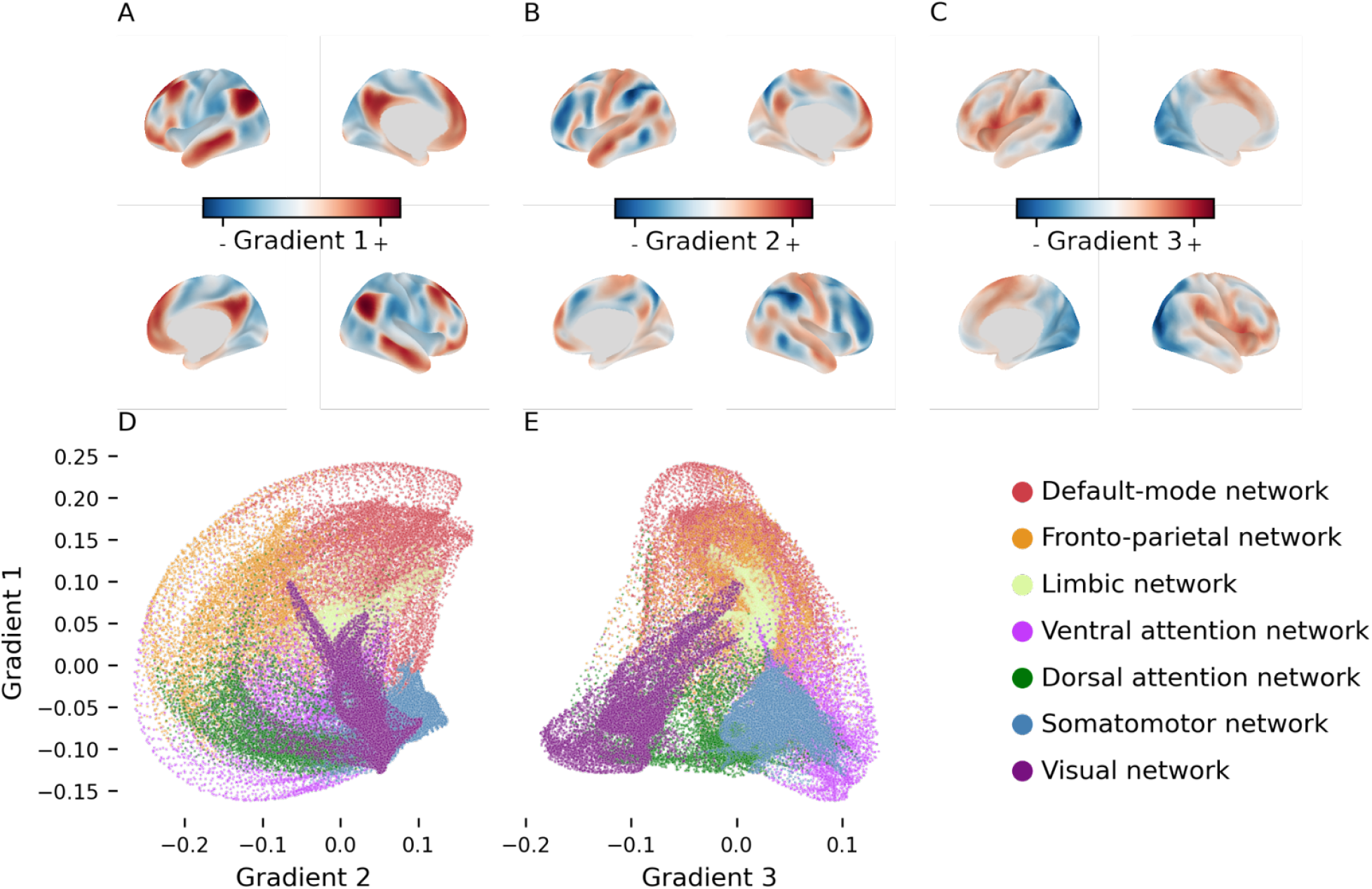
**A-C**: The first three gradients of functional connectivity (group median) displayed on the inflated cortical surface. These three dimensions respectively recapitulate a different functional axis each: sensory-DMN axis (**A**), FPN-DMN (**B**), and visual-somatomotor (**C**). **D-E**: scatterplot of the median location of all cortical vertices along the first and second gradient (**D**), and first and third gradient (**E**). Vertices are colored based on their affiliation to the seven resting-state functional networks as per legend (bottom right).

In the present work, we investigate interindividual FC variability within the functional gradient framework to better understand how variability relates to different facets of intelligence. We used generalized canonical correlation analysis (GCCA) to decompose the resting-state BOLD time series of 338 unrelated subjects from the Human Connectome Project (HCP), Young Adults dataset (11). We identified the three first components that best explain the activity patterns across subjects. These gradients effectively describe the activation profile of any given point on the cortical surface by locating it in a tridimensional space where its coordinates (gradients) represent its affinity with different functional systems. Thus, to map FC variability across the cortex, we estimated the dispersion (12) in *gradient space* of each vertex on the cortical surface sampled from every individual. Eight clusters were identified from this map where dispersion is maximal and separately tested the association between the three gradients at these loci and intelligence measures. This approach allowed for testing the association of functional variability along separate, interpretable dimensions with behavior. Specifically, we studied the effect of each of the clusters’ gradients on individual scores of fluid intelligence, crystallized intelligence, and general intelligence (the g-factor). Then, to further understand the implications of gradient dispersion, we assessed whether gradients relevant to cognitive scores are also associated with variability of topological properties of FC profiles.

## 2. Results

### 2.1 Interindividual variability of functional gradients

The three dimensions revealed by the GCCA had a spatial distribution (Fig. 1) in line with the gradients previously described by Margulies and colleagues (9). However, the order of the second and third gradient was inverted. In other words, in our analyses the FPN-DMN gradient explained more variance in the blood oxygenation level dependent (BOLD) signal time series than the visual-somatomotor one.

Functional gradients capture the latent dimensions that guide the covariance patterns of BOLD signal across the cortex. Consequently, cortical regions are arranged along each gradient based on the similarity of their patterns of activity with respect to a specific functional dimension. Together, they define an n-dimensional space where spatial distance between regions reflects their functional distance. Interindividual variability of FC was measured by identifying the group-level centroid of each vertex in gradient space, and computing the sum of the squared distance from individual vertices to the centroid (12). The resulting cortical map of gradient *dispersion* (Fig. 2A) showed a pattern in line with previous studies (13), with peaks of interindividual variability in the dorso-lateral temporal cortex extending to the inferior temporal lobule, and in the ventro-lateral, dorso-lateral, and anterior-medial frontal cortex (Fig. 2B). Additionally, a variability locus emerged in the occipital pole that had not been described before (Fig. 2B).

**Figure 2.**
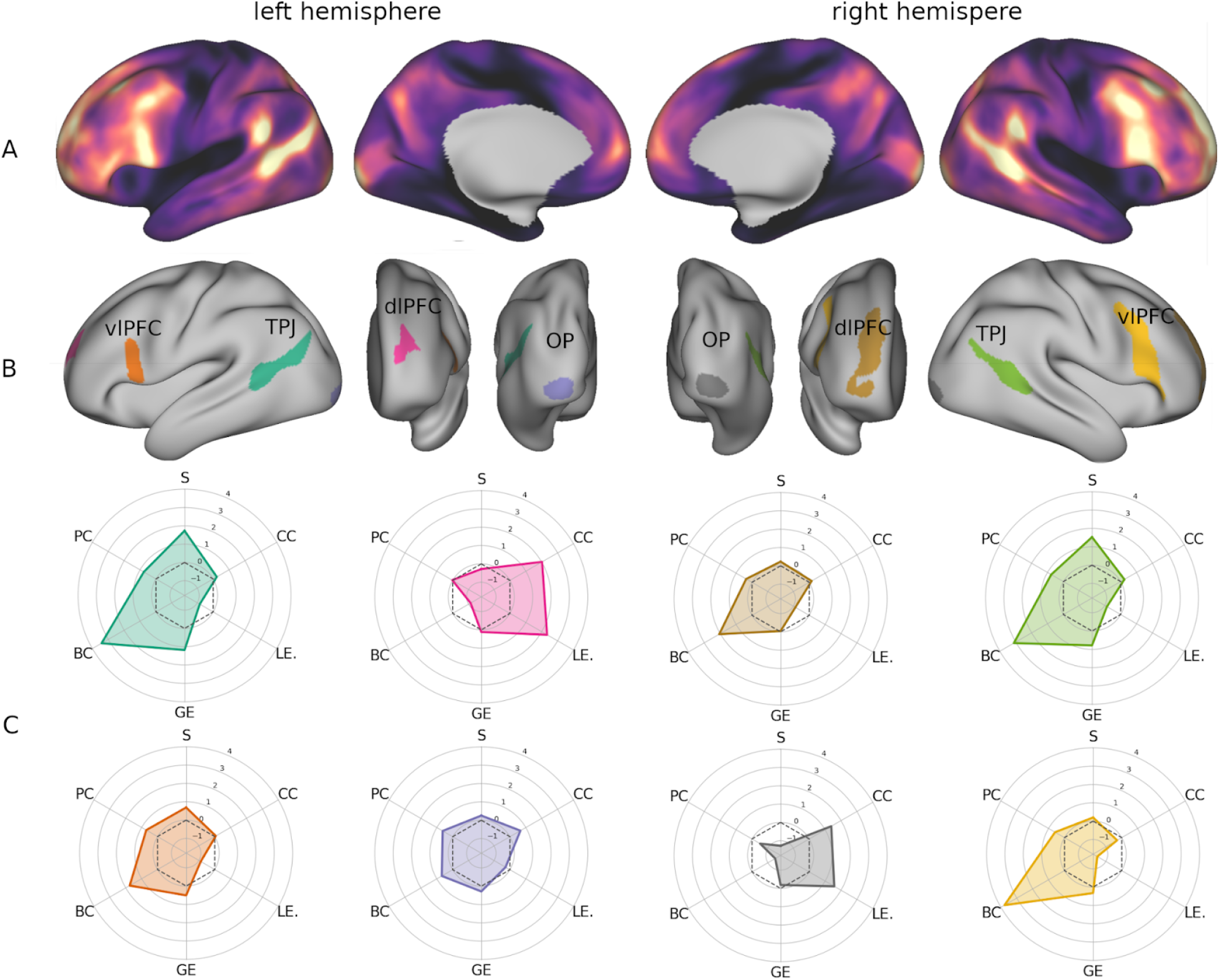
**A** Map of cross-subject vertex dispersion in gradient space displayed on the inflated cortical surface. **B** Clusters of high interindividual variability obtained by thresholding the surface map of interquartile range at the 95^th^ percentile. **C** Topological profiles of the variability clusters visualized as radar plots. The plots are colored based on the cluster they correspond to in panel B. TPJ: temporo-parietal junction; vlPFC: ventrolateral prefrontal cortex; dlPFC: dorsolateral prefrontal cortex; OP: occipital pole; S: strength; CC: clustering coefficient; LE: local efficiency; GE: global efficiency; BC: betweenness centrality; PC: participation coefficient.

To understand how much variability along each gradient dimension contributed to global dispersion of FC, we tested its correlation with dispersion measured along each individual component separately.

Dispersion of the vertices’ FC profiles appeared more closely related to the variability on the FPN-DMN axis (ρ = 0.95, p < 0.001), followed by the sensory-DMN axis (ρ = 0.85, p < 0.001), and the visual-somatomotor axis (ρ = 0.60, p < 0.001). Moreover, dispersion was also positively correlated with the principal gradient (ρ = 0.34, p < 0.001) and negatively correlated with the other two functional axes (FPN-DMN: ρ = −0.33, p < 0.001; visual-somatomotor: ρ = −0.12, p < 0.001), indicating that variability was higher in the vertices near the DMN, FPN, and visual ends of the three gradients respectively.

### 2.2 Clusters of maximum variability

In order to identify the loci where gradient variability across subjects is maximal, we thresholded the map shown in Fig. 2A at the 95^th^ percentile. From this process eight clusters of maximum interindividual variability (Fig. 2B) emerged, four in each hemisphere. Two of them were located in the bilateral temporo-parietal junction (TPJ) and they overlapped with DMN (L: 19%, R: 30%), VAN (L: 41%, R: 34%), and DAN (L: 40%, R: 36%). Two more clusters occupied the occipital poles and were entirely enclosed in the visual network. The remaining four clusters were found in the dorsolateral and ventrolateral prefrontal cortex (dlPFC, vlPFC): two in the rostral middle frontal sulcus, and two in the opercular portion of the inferior frontal gyrus extending to the inferior frontal sulcus and middle frontal gyrus. Despite a similar placement, the right prefrontal clusters were far more extended and overlapped with different networks compared to their left counterparts. Most vertices of the left vlPFC cluster belonged to VAN (58%), with smaller portions affiliated to FPN (21%), DAN (18%), and DMN (3%). The right vlPFC cluster, instead, intersected mainly with FPN (59%), sharing fewer vertices with DMN (26%), VAN (14%), and DAN (1%). Lastly, the dlPFC cluster overlapped almost equally with FPN (51%) and DMN (46%) in the right hemisphere, but was almost entirely included within DMN (75%) in the left, with smaller portions intersecting VAN (16%) and FPN (9%).

### 2.3 Associations between functional gradients and intelligence

After identifying the loci where interindividual FC variability was maximal, we aimed to investigate its relationship with cognitive capabilities. Thus, we examined how the location of these loci along each functional gradient related to fluid, crystallized, and general intelligence (see Methods for further detail on how these measures were obtained). To this end, a multiple linear model was tested for each intelligence measure and each functional dimension using the clusters’ gradients as predictors, controlling for age, handedness, sex, and education. The gradients of the TPJ and occipital clusters were highly correlated between hemispheres (TPJ: first gradient, ρ = 0.66; second gradient, ρ = 0.61; third gradient, ρ = 0.61; occipital pole: first gradient, ρ = 0.94; second gradient, ρ = 0.96; third gradient, ρ = 0.97). To avoid including redundant explanatory variables to the models, the left and right homologues of these clusters were averaged and included as a single predictor each.

The only model that survived significance testing was the prediction of fluid intelligence based on the principal gradient of the variability clusters (F = 2.88, p = 0.012). This result was driven by a significant negative effect of the principal gradient of the right vlPFC cluster (β = −57.16, t = −2.62, p = 0.008; Fig. 3G). To obtain a more cognitively detailed understanding of this association, we then tested the correlation between the principal gradient of this cluster and individual scores of the tasks included in the fluid intelligence measure using Spearman’s correlation coefficient. Weak but significant associations were found with the dimensional change card sorting task (ρ= −0.15, p = 0.023; Fig. 3H) and the flanker inhibitory control and attention task (ρ= −0.19, p = 0.002; Fig. 3I). Overall, these results indicate that when the right vlPFC is functionally further from the DMN and closer to sensory systems (i.e. lower principal gradient values), individuals perform better in tasks testing executive control.

**Figure 3.**
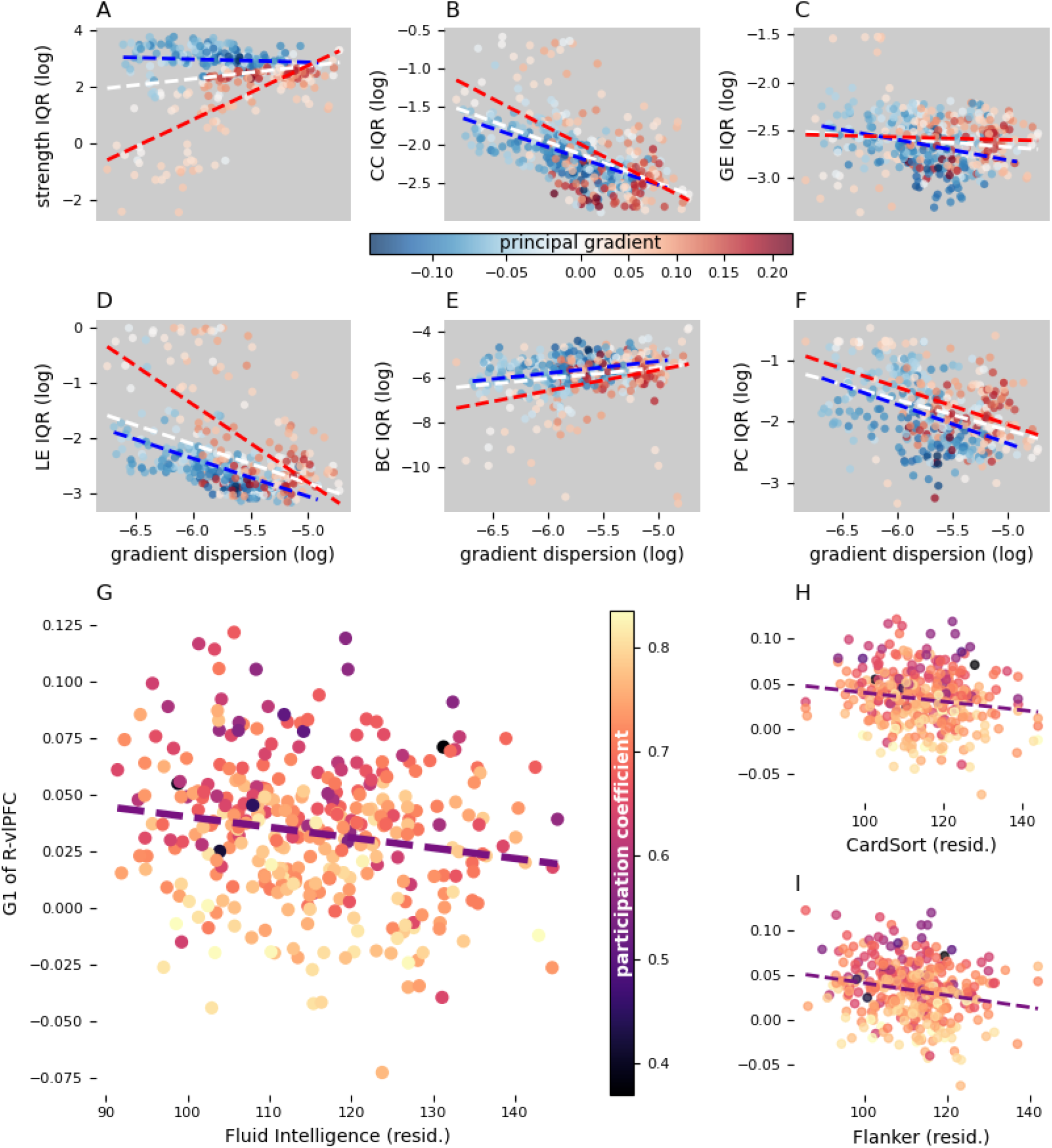
**A-F** scatterplots of the interindividual variability of graph metrics against gradient dispersion (**A**: strength; **B**: clustering coefficient (CC); **C**: global efficiency (GE); **D**: local efficiency (LE), **E**: betweenness centrality (BC); **F**: participation coefficient (PC)). Each marker represents a graph node. Marker color represents the principal gradient (group median) of the corresponding node. Over the scatter plot are the regression lines (dashed lines) of the nodes with a negative gradient (blue), positive gradient (red), and all nodes together (white). **G-I** Fluid intelligence (**G**), card sorting (**H**), and flanker inhibition (**I**) scores plotted against the principal gradient of the right ventrolateral prefrontal cluster after correction for age, gender, education, and handedness. Each marker represents the principal gradient and cognitive scores of an individual and its color represents the cluster’s participation coefficient. G1: gradient 1; resid.: residuals; R-vlPFC: right ventrolateral prefrontal cortex; IQR: interquartile range.

### 2.4 Topological properties of the variability clusters

The final set of analyses aimed to explore the relationship between the interindividual variability of functional gradients and topological properties of cortical regions. To this end, the Schaefer atlas (400 parcels) was modified to include the eight variability clusters eroding the original ROIs where they overlapped with them. Graph theory tools were employed for the construction of individual FC graphs, in which regions are treated as nodes and their functional correlation as edges connecting them. To limit the risk of Type 1 error, graphs were thresholded preserving only the top 10% of all the connections, in line with prior large cohort studies proving that 10% sparsity threshold have the highest test-retest reproducibility (14). The *topological profile* of each parcel was then drawn by calculating: the total strength of connections, local efficiency, clustering, global efficiency, betweenness centrality, and participation to multiple intrinsic functional networks.

As can be qualitatively appreciated in Fig. 2C, the topological profiling revealed that the right vlPFC was, at the same time, the cluster with the highest betweenness centrality (z = 3.78) and lowest local efficiency (z = −1.54) and clustering (z= −0.29). Other clusters showed a similar pattern but not as pronounced. For example, the temporal clusters did show a high centrality (L: z = 3.42, R: z = 3.11), however their local efficiency was not as markedly low (L: z= −0.85, R: z= −0.87). The topological properties of the right vlPFC suggest that it may have a role in mediating communication between functionally segregated systems.

Expecting regions with highest gradient variability to also have an inhomogeneous topological profile across individuals, we assessed whether the group-level cortical maps of gradient dispersion and graph metric variability (interquartile range) correlated significantly. Contrary to our expectations, however, the across-individual interquartile range of many graph metrics correlated negatively with gradient variability (clustering: ρ = −0.60, p < 0.001; local efficiency: ρ = −0.48, p < 0.001; participation coefficient: ρ = −0.39, p < 0.001). The only exceptions were the variability map of betweenness centrality, which showed a positive correlation with gradient dispersion (ρ = 0.23, p < 0.001), and global efficiency, which did not show any significant association. Interestingly, the scatterplot of strength variability against gradient dispersion suggested the existence of two sub-population of parcels with an opposite association between the two measures. Indeed, further testing revealed that parcels closer to the sensory end of the principal gradient show a negative correlation between the variability of strength and gradients (ρ = −0.36, p < 0.001), while those closer to the DMN end display a positive association (ρ = 0.60, p < 0.001).

Finally, we investigated how topological properties related to the principal gradient, with a specific focus on the right vlPFC cluster. Thus, we measured the across-subject correlation between each graph metric and principal gradient for every parcel. In the right vlPFC, a significant negative correlation emerged between the participation coefficient and principal gradient (ρ = −0.53, p < 0.001), suggesting that, when this region is functionally closer to sensory and attentional systems, its connections are more evenly distributed across a larger number of networks. No other topological property of this cluster varied together with the principal gradient across individuals.

A detailed description of the correlation maps is beyond the scope of this paper, however it is worth noting that direction and magnitude of the correlation coefficients were modulated by the principal gradient. For instance, parcels on the sensory half of the gradient display higher strength and global efficiency when they are closer to the lower end of the axis, but this association is reversed in the opposite half of the gradient.

## 3. Discussion

This study demonstrates that the FPN’s placement between sensory processing and abstract cognition systems relates to intelligence, and shows how graph topology can help interpret this relationship. The spatial distribution of cross-individual gradient dispersion is analogous to that of FC variability measured through traditional connectivity analyses, with interindividual differences peaking in TPJ, vlPFC, and dlPFC (4). An additional locus of high variability, however, was also found in the occipital pole. Regression modeling revealed that individual fluid intelligence scores are associated with the coordinate of the right vlPFC along the functional axis going from sensory areas to the DMN. Graph-theory analyses, suggests that this factor also contributes to this region’s ability to participate in multiple functional systems.

The pattern of gradient variability captured by our analyses is in line with previous publications indicating associative regions as the most variable across individuals throughout multiple imaging modalities (4, 7, 8, 13). This is likely a consequence of their prolonged postnatal development (15, 16), which makes their connectivity and morphology less subject to genetics and more susceptible to variable environmental influence (5, 17). For instance, sensory projections differ less between individuals compared to long-range fiber tracts (18), which support long-distance FC distinctive of association areas (19). The differences in long-range connectivity are thought to also underlie the high variability of cortical folding patterns that characterizes association regions (20, 21). Recent studies, however, demonstrated that cortical topography and spatial relationships also have a strong influence on functional organization (22, 23). In particular, a large portion of the variability of individual FC profiles is attributable to differences in spatial arrangement and topographical overlap of cortical parcels (24, 25). The principal functional gradient itself shows a clear spatial organization as sensory regions are located furthest from DMN (9, 26). It cannot be ruled out, therefore, that part of the FC variability observed here and in previous studies (8) is due to topographical differences across individuals.

Functional gradients allow a neurocognitive interpretation of functional variability, combining a topological and a cognitive interpretation of our findings. The principal gradient (sensory-DMN) was the only neurocognitive dimension to significantly relate to intelligence, despite tridimensional dispersion being more closely related to variability along the DMN-FPN axis. Given that the sensory-DMN axis explains the most variance in intrinsic activity (9, 10), it is conceivable that interindividual variability along this dimension would more affect individual cognitive abilities. In line with existing literature, such variability was specifically associated with fluid intelligence and executive control. This relationship was driven by a negative association between these cognitive measures and the principal gradient of the right vlPFC. That is, a larger functional segregation between this region and the DMN benefits executive function. A possible interpretation of this effect is that moving further from the DMN, the vlPFC acquires an intermediate topological position on the principal gradient between this network and sensory/attention systems, thus acting as a junction between them. This hypothesis was corroborated by the topological profiling of the cluster, which exhibits, on average, high betweenness and low local efficiency. That is, the vlPFC is traversed by the shortest paths between many node pairs, but it is linked to regions that are loosely connected with one another (27). Thus, this region may act as a connector hub coordinating otherwise highly segregated systems, likely the sensory and DMN ends of the principal gradient. In line with this interpretation, further analyses confirmed that the connectivity profile of right vlPFC is more evenly distributed across multiple networks when it is located closer to the sensory end of the principal gradient. Considering the influence that cortical geometry has on FC measures (24, 25), this result could also indicate that a more median placement on the principal gradient is associated with being evenly distanced from all systems. Either these properties may be instrumental in guiding efficient interaction and interchange between externally- and internally-driven processing: a key factor in successful executive control. Multiple studies have shown that during tasks probing executive function both attentional systems and DMN are alternately recruited based on which of these two processing modalities is required at any given time (28–32). These publications used protocols very similar to the card sorting and flanker tasks used in the HCP dataset, which are the components of fluid intelligence that most closely relate to principal gradient. Hence, the placement (topological or topographycal) of the right vlPFC between modality-specific and transmodal systems may interact with this specific aspect of executive function.

In conclusion, this study expands our current knowledge on interindividual variability of the functional architecture of the cortex by decomposing it into interpretable neurocognitive axes intrinsic to individual functional patterns. Our results suggest that the link between individual functional fingerprint and cognitive performance is driven by the ability of prefrontal cortices to coordinate internal schemas in response to external sensory input.

## 4. Methods

### 4.1 Participants & Data

The study was based on openly accessible data provided by the WU-MInn HCP initiative (11). The sample included 338 healthy, unrelated participants (F=188, M=158) of 28.6 years of age on average (SD = 3.6). For all subjects, four rfMRI time series of about 15 minutes were available, as well as scores at several off-scan cognitive tasks including measures of memory, attention, and executive functions. The rfMRI time series were acquired through a 3-T Siemens connectome-Skyrausing a gradient-echo EPI sequence with 2 mm isotropic voxels, 720 ms TR, and 52° flip angle. The HCP minimal preprocessing pipeline for these data is described in detail in (33) and includes correction for spatial distortion, motion, and bias field, and registration to T1w image. Temporal artifacts were cleaned from the data using high-pass filtering and independent component analysis (34). The voxels’ time series were mapped to the native surface and registered to the 32k Conte69 mesh applying 2 mm FWHM smoothing. This process ensures spatial correspondence of vertices across subjects. To account for residual anatomical variability, we applied additional smoothing of 6 mm FWHM to the functional time series when computing vertex-wise GCCA. Smoothing was also performed prior to averaging rfMRI time series within parcels to compute FC in order to ensure consistency of the data analyzed across methodological approaches.

### 4.2 Functional connectivity gradients

Functional gradients are latent dimensions of FC space using different dimensionality-reduction algorithms (10) along which the cortical vertices are placed based on the similarity of their connectivity profiles with respect to that specific dimension. For example, vertices within the visual and somatosensory systems will be very close on the sensory-DMN axis, but very distant on the visual-somatosensory axis (Fig. 1D-E). To derive these latent dimensions we applied generalized canonical correlation analysis (GCCA) (35, 36) (36) to the four BOLD time series that had previously been normalized and concatenated. GCCA generalizes principal component analysis (PCA) across more than two data sets by finding the latent factors that maximize pairwise correlations between all sets. The specific algorithm used here initially computes the most informative principal components of each subject’s BOLD time series and then applies singular value decomposition to a concatenated version of these components ([mvlearn.embed.GCCA](https://mvlearn.github.io/references/embed.html#generalized-canonical-correlation-analysis-gcca)). This second decomposition is used to compute projection matrices that project the subjects’ data into highly correlated subspaces, whose dimensions represent the functional gradients. Our analysis was limited to the extraction of the three gradients that explained the most variance.

### 4.3 Interindividual variability of functional connectivity profiles

Interindividual variability of FC profiles was estimated by measuring the dispersion of vertices in the tridimensional space defined by functional gradients. Specifically, we adopted the procedure proposed by Bethlehem and collaborators (12) to capture between-subjects variability. To this end, a group-level centroid was identified for each vertex based on its coordinates on the three gradients in across subjects. The dispersion around this point was then measured as the sum squared Euclidean distance of the individual vertices from it (Fig. 2A). The same approach was also applied to determine vertex dispersion along each gradient’s axis. Loci of maximum variability were defined by thresholding the functional dispersion map at the 95^th^ percentile and applying a 200 mm^2^ cluster-size threshold, which resulted in the identification of 8 separate clusters (Fig. 2B).

### 4.4 Modeling intelligence

Three intelligence scores were computed for all subjects: crystallized intelligence, fluid intelligence, and g-factor. These measures were calculated by averaging multiple cognitive measures of the HCP off-scan battery. Crystallized intelligence included the picture vocabulary and the oral reading recognition tests; fluid intelligence included the dimensional change card sorting test, the flanker inhibitory control and attention test, the picture sequence memory test, the list sorting working memory test, and Penn progressive matrices; the g-factor included all the measures above, the pattern comparison processing speed test, variable short Penn line orientation test, and the Penn word memory test. While the first two measures were used as provided by the HCP, the g-factor was computed as the mean of its components weighted by their loadings in the factor analysis performed by Dubois and colleagues (37, 38) (see Table 1). To control for confounding variables, the effects of age, handedness, sex, and years of education were regressed out from both intelligence and gradient measures.

**Table 1.** Subset of task scores used to estimate crystallized intelligence, fluid intelligence, and the *g* factor for all subjects. The task weights used for computing the *g* factor are in accordance with prior literature (Dubois et al., 2018; Lohmann et al., 2021).

| Cognitive component |  | Test | Cognitive construct | Weight |
| --- | --- | --- | --- | --- |
| g-factor | Crystallized intelligence | Picture vocabulary | Language/vocabulary comprehension | 0.624 |
|  |  | Oral reading recognition | Language/reading decoding | 0.642 |
|  | Fluid intelligence | Dimensional change card sorting | Executive function/cognitive flexibility | 0.364 |
|  |  | Flanker inhibitory control and attention task | Executive function/inhibition | 0.259 |
|  |  | List sorting | Working memory | 0.451 |
|  |  | Penn progressive matrices | Fluid intelligence | 0.626 |
|  |  | Picture sequence Memory | Episodic memory | 0.354 |
|  | – | Pattern comparison | Processing speed | 0.232 |
|  |  | Variable short Penn line orientation | Spatial orientation | 0.578 |
|  |  | Penn word memory test | Verbal episodic memory | 0.294 |

For each intelligence measure three multiple linear models were built using respectively the first, second, and third gradient of the maximum-variability clusters as explanatory variables. The significance of this model was tested through a permutation approach by comparing the F-value of the original model against a null F distribution obtained by permuting the dependent variable 10,000 times. The p-value was calculated as the fraction of null F values higher than the original one. Post-hoc tests were performed by applying the same procedure to the t-values of the individual predictors. In this case, the p-value was calculated on the distribution of absolute t-values. All p-values were corrected for false discovery rate (FDR) of 0.05.

### 4.5 Variability clusters in network topology

The subjects’ cortical surface was divided into 400 parcels according to the atlas by Schaefer et al. (2018), and the percentage of overlapping vertices between the clusters and resting state networks was calculated. We then added the clusters defined by our analyses to the Schaefer atlas masking the original parcels where the two overlapped, so that downstream analyses would only consider parcel vertices that were not shared with any other ROI. We obtained a total of 406 parcels (L: 204, R: 202), as two of the original parcels were entirely masked by the right TPJ and vlPFC clusters. To build functional connectivity graphs, a 406×406 adjacency matrix was computed for every individual using the Pearson’s correlation coefficient between parcel-averaged BOLD time series. All self connections were removed from the adjacency matrices which were then thresholded preserving only the 10% strongest positive edges in the graph to reduce the risk of including false positive connections. From these networks, six topological metrics of our clusters were calculated: i) strength (total strength of all connections); ii) global efficiency (average closeness to all the other nodes); iii) local efficiency (average global efficiency within a node’s neighborhood); iv) clustering (the portion of a node’ neighbors also connected with each other); v) betweenness centrality (the number of shortest paths that pass through a node); vi) participation (how distributed a node’s edges are across all communities in the graph). To calculate the latter, seven communities were defined *a priori* based on the seven canonical networks (39) defined by Schaefer and colleagues (40). Since the variability clusters overlapped multiple networks, they were assigned to a separate community.

## Acknowledgements

Data were provided by the Human Connectome Project, WU-Minn Consortium (Principal Investigators: David Van Essen and Kamil Ugurbil; 1U54MH091657) funded by the 16 NIH Institutes and Centers that support the NIH Blueprint for Neuroscience Research; and by the McDonnell Center for Systems Neuroscience at Washington University.

